# Dissecting the Tumor Microenvironment at Single-Cell Resolution to Prioritize Candidate Tumor Antigens Across Multiple Cancer Types

**DOI:** 10.64898/2026.11.24.768856

**Authors:** Thayana Vieira Tavares, Pedro Lucas Alves da Silva, Laize Barros Pantoja, Jorge Estefano Santana de Souza, André Luis Fonseca Faustino

## Abstract

Cancer immunotherapy has emerged as a revolutionary therapeutic approach, advancing the treatment of several cancer types. However, the identification of candidate tumor antigens remains challenging, as multiple lines of evidence are required to establish malignant-cell enrichment and therapeutic suitability. This study aimed to characterize and prioritize candidate tumor antigen genes based on their enrichment in malignant cells through the integration of single-cell transcriptomic data with complementary immunological and clinical evidence from the literature and public databases. An integrated pan-cancer dataset comprising 355,941 cells across 10 tumor types and 21 cell populations was analyzed using enrichment categories adapted from the Human Protein Atlas (HPA). Candidates were characterized according to antigen classification, gene expression in healthy tissues, presence of reported epitopes, recurrence across tumor types, and associations between gene expression and overall survival. Among the 22,779 genes evaluated, 30 candidates were retained after the prioritization process. CLDN6, MAGEC2, and PLAC1 stood out due to previous investigations in recent clinical immunotherapy research. Additionally, 16 of these 30 candidates showed significant associations between gene expression and overall survival across distinct cancer types, potentially supporting their prioritization for further investigation as immunotherapy targets. In contrast, some cancer-testis antigens (CTAs), previously regarded as promising therapeutic targets, were found to have limited suitability due to their enriched, enhanced, or group-enriched expression in non-malignant cells. Most of these steps can be independently reproduced using the publicly available Nextflow pipeline developed for this study.

## 1 Introduction

Cancer immunotherapy has emerged as a revolutionary therapeutic approach, advancing the treatment of several types of cancer compared to existing therapeutic strategies, such as chemotherapy and radiation therapy (Zhang; Zhang, 2020; Ahuja; Zaheer, 2024). There are several types of cancer immunotherapy in development, including T cell therapies, CAR-T cells, cancer vaccines, and monoclonal antibodies, all these approaches rely on the identification of antigens to effectively target malignant cells and elicit anti-tumor immune responses (Kuznetsova et al., 2024).

Bulk RNA sequencing (RNA-seq) has been widely used to identify candidate antigen genes differentially expressed in tumor samples, reflecting its widespread adoption as an established technology in cancer research (Li and Wang, 2021). However, this approach provides an average representation of gene expression in tumor microenvironment, rather than distinguishing malignant cells from other cellular populations, tumors contain diverse immune and stromal populations, including T cells, B cells, natural killer (NK) cells, myeloid cells, fibroblasts, and endothelial cells. Their composition and cellular states vary across tumor types and among regions of the same tumor (Giraldo et al., 2019; Wu et al., 2019). Furthermore, gene expression does not necessarily guarantee protein production. Single-cell proteomics represents a recent technology that could potentially address both limitations by providing detailed insights into tumor microenvironment heterogeneity and the presence of molecules at the protein level. However, its application remains challenging due to limitations in sensitivity and proteome coverage, in addition to its high cost (Zeng; Wang, 2023).

To bridge these gaps, the integration of single-cell RNA-seq (scRNAseq) data with complementary lines of evidence represents a strategic approach. ScRNAseq enables gene expression to be characterized at cellular resolution, allowing malignant cell expression to be distinguished from non-malignant components and intratumoral heterogeneity to be investigated (Ren et al., 2021). This resolution has also been applied to the discovery of targets for cellular therapies (Huang et al., 2023). In addition to scRNAseq, other lines of evidence are needed to identify promising candidates for potential tumor antigens based on the heterogeneity of the microenvironment, including the investigation of peptide evidence and the analysis of gene expression in healthy tissues and non-malignant cells of the tumor microenvironment (Haen et al., 2020). The enrichment of antigens in non-malignant cells and healthy tissues can compromise target selectivity and lead to on-target, off-tumor toxicity, what is the toxicity resulting from target recognition in non-tumor tissues (Zhang et al., 2021).

Tumor antigens can be broadly classified by origin and expression pattern as tumor-associated antigens (TAAs) or tumor-specific antigens (TSAs). TAAs derive from proteins that are also expressed, to varying degrees, in normal tissues, but are enriched in malignant cells, whereas TSAs are restricted to tumor cells and include neoantigens generated by tumor-specific genomic alterations, such as somatic mutations (Schumacher and Schreiber, 2015). Categories of shared TAAs include differentiation-associated antigens, overexpressed antigens, and cancer-testis antigens (CTAs) (Buonaguro et al., 2010). Differentiation-associated antigens are linked to specific cell lineages and can be expressed by both tumor cells and their normal tissue of origin. Overexpressed antigens are also present in normal tissues but occur at higher levels in tumor cells (Alarcon et al., 2022; Buonaguro et al., 2010).

As for CTAs, they constitute a distinct subset of TAAs characterized by expression in multiple tumor types and physiological expression largely restricted to germline cells, particularly the testis, with some members also expressed in the placenta (Scanlan et al., 2002). These characteristics have made CTAs attractive targets for immunotherapeutic approaches, including cancer vaccines and TCR-based therapies (Fan et al., 2021). However, for a gene to be considered an acceptable immunotherapeutic target, it should also exhibit low or no expression in healthy tissues other than germline tissues, as well as low expression in non-malignant cells (Haen et al., 2020). In this regard, identifying CTAs with high enrichment in non-malignant cells can help exclude candidates with lower therapeutic potential, whereas their enrichment in malignant cells can highlight genes with greater potential as immunotherapeutic targets.

This study aimed to characterize and prioritize candidate tumor antigen genes based on their enrichment in malignant cells through the integration of single-cell transcriptomic data with complementary immunological and clinical evidence from the literature and public databases. This approach enabled the investigation of the cellular enrichment patterns of all genes available in the dataset, including CTAs in non-malignant cells, the identification of candidates shared across tumor types, the assessment of reported epitope evidence in public databases, the evaluation of prioritized candidates expression in healthy tissues, and the analysis of associations between candidate gene expression and patient overall survival using publicly available bulk RNAseq data. Additionally, the literature was reviewed to identify immunotherapeutic strategies under investigation targeting genes prioritized in this study across different cancer types.

## 2 Materials and Methods

### 2.1 Nextflow pipeline developed for this study

A Nextflow DSL2 pipeline was developed to automate and ensure the reproducibility of the main analyses presented in this work. Although originally designed for these analyses, the pipeline can also be adapted to other datasets, supporting its reuse in independent studies. It integrates R 4.4.1 and Python 3.13 scripts into two complementary workflows.

The first workflow processes AnnData objects in H5AD format, generating an interactive dataset summary and identifying genes enriched in malignant cells and other cell populations. Candidates are further characterized by antigen type, expression across non-malignant cells and healthy tissues based in HPA data with corresponding visual summaries. The second workflow integrates epitope evidence provided by the user, compares candidate gene expression patterns across malignant cells from different tumor types, when multiple cancer data is available in the H5AD file, performs functional enrichment analysis, and prioritizes candidates based on malignant cell enrichment, HPA tissue specificity, epitope evidence reported, antigen classification, and pan-cancer recurrence. The pipeline is publicly available on GitHub at https://github.com/LIIA-edu/tumor-antigen-pipeline.

### 2.2 Data collection

The data were obtained from Gondal, Cieslik, and Chinnaiyan (2025) and comprised integrated scRNAseq datasets from eight independent studies. The atlas included 10 annotated tumor types, nine of which contained malignant cells: triple-negative, estrogen receptor-positive, and HER2-positive breast carcinomas, basal cell carcinoma, melanoma, metastatic melanoma, nonpapillary renal cell carcinoma, intrahepatic cholangiocarcinoma, and hepato-cellular carcinoma. Squamous cell carcinoma was the only tumor type without annotated malignant cells. The integrated dataset included 90,270 malignant cells and 355,941 non-malignant cells. The original authors performed quality control, preprocessing, and integration; the resulting data were downloaded in H5AD format from the CZ CELLxGENE Discover platform.

### 2.3 Merge similar cell types and remove unknown cells

Cell-type annotations were retained as reported by the original authors, however, cells annotated as Unknown were excluded before downstream analyses so that analyses were restricted to classified cell populations. Related T-cell subtypes were grouped to facilitate downstream analyses focused on malignant cells rather than T-cell heterogeneity. The CD4 T-cell category included CD4-positive alpha-beta T-cells, T follicular helper cells, T-helper 17 cells, and regulatory T-cells. The CD8 T-cell category comprised CD8-positive alpha-beta T-cells, activated and effector CD8-positive alpha-beta T-cells, central memory CD8-positive alpha-beta T-cells, naive thymus-derived CD8-positive alpha-beta T-cells, and CD8-positive CD28-negative alpha-beta regulatory T-cells. The dataset also included generic T-cell annotations, such as T-cells and naive T-cells.

### 2.4 Malignant cell definition

Malignant-cell copy-number variation (CNV) classifications were retained as defined in the original studies. HoneyBADGER was used in the Pozniak et al. (2024) and Yost et al. (2019) cohorts, and inferCNV was used in the Jerby-Arnon et al. (2018), Tirosh et al. (2016), Bassez et al. (2021), Bi et al. (2021), and Ma et al. (2019) studies. In Alvarez-Breckenridge et al. (2022), malignant cells were identified from inferred copy-number profiles in the CD45^*−*^, CD45^+^, and CD3^+^ categories.

### 2.5 Detection rate as a cell type enrichment metric

The detection rate was defined as the proportion of cells within a given cell type in which a gene was detected. A gene was considered detected in a cell when its expression value was greater than zero in the processed expression matrix.

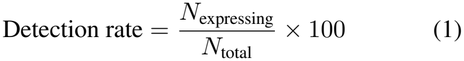

### 2.6 Adaptation of Human Protein Atlas (HPA) enrichment definition

Cell-type enrichment metric was assessed using a scheme adapted from the Human Protein Atlas (HPA), which classifies genes by enrichment across biological contexts, including tissues, brain regions, cell lines, immune cell types, and single-cell populations. The five categories used were Enriched, Group Enriched, Enhanced, Low Specificity, and Not Detected. Genes detected in fewer than 1% of cells were classified as Not Detected; genes detected in at least 1% of cells but not matching another enrichment criterion were classified as Low Specificity. Enriched genes were expressed in one cell type at least four times more than in any other cell type. Group Enriched genes were enriched across a group of two to five cell types. For this study, Enhanced was defined as expression in the target cell type at least four times greater than the mean expression in all other cell types, excluding the target cell type from that calculation. Genes enriched in malignant cells were considered the initial candidate antigen genes.

### 2.7 Classification of antigen-related gene categories

Candidate antigen genes were classified according to antigen categories curated from the Tumor T-cell Antigen Database (TANTIGEN) (Olsen et al., 2017) and the study by Da Silva et al. (2017). The categories included Differentiation, Overexpressed, and Cancer-Testis antigens, genes not assigned to any of these three categories were labeled as unclassified. CTAs classified as Enriched, Group Enriched, or Enhanced in non-malignant cell populations were further investigated using four HPA RNA datasets: single-cell, single-cell tissue, single-nucleus brain, and immune-cell expression datasets to assess their expression in healthy tissues. Furthermore, the initial candidate antigen genes were investigated using publicly available databases, including Gene Ontology platform (QuickGO) (Binns et al., 2009), Oncology Knowledge Base (OncoKB) (Chakravarty et al., 2017), and Cell Surface Protein Atlas (CSPA) (Bausch-Fluck et al., 2015), to retrieve additional protein information.

### 2.8 HPA-based specificity assessment of candidate antigen genes

Candidate antigen genes were queried against the Human Protein Atlas (HPA) API to retrieve RNA expression specificity and distribution data across tissues and healthy single-cell populations. The retrieved information included tissue-level and single-cell RNA specificity classifications and expression values (nTPM and nCPM).

### 2.9 Curation of epitopes distribution of candidate antigens

An automated Python script (Python 3.12) using Play-wright (version 1.62.0) retrieved epitope information from the Immune Epitope Database (IEDB) (Vita et al., 2018) and the Cancer Epitope Database and Analysis Resource (CEDAR) (Koşaloğlu-Yalçın et al., 2022), using the search options available on each platform. For both databases, the search was restricted to linear epitopes associated with multiple cancer types specified in the database queries, T-cell or MHC ligand assays, MHC class I, and human hosts. CEDAR records were additionally filtered by epitope origin to retain germline-derived, self-derived, or host-derived antigens.

### 2.10 Curation of tumor-specific distribution of candidate antigen genes

To assess tumor-specific distribution, candidate antigens were evaluated independently in malignant cells from each of the nine tumor types in the integrated dataset. A gene was considered present in a tumor type when detected in at least 1% of its malignant cells. These data were used to construct a binary gene-by-tumor presence-absence matrix, quantify tumor-specific and shared candidates, and visualize overlap with an upset plot.

### 2.11 Literature review of candidate antigen genes in investigational immunotherapies

Candidate antigen genes were investigated in the scientific literature to identify previous or ongoing research exploring their use in immunotherapeutic approaches, including cancer vaccines, T-cell-based therapies, and other immunotherapy strategies. Searches were conducted using scientific literature platforms, as well as clinical trial registries and research platforms, such as ClinicalTrials.gov (Zarin et al., 2011), to identify studies evaluating these genes as potential immunotherapeutic targets.

### 2.12 Prioritization candidate antigen genes

Candidate antigen genes were prioritized by integrating multiple sources of evidence, including tissue expression specificity from the Human Protein Atlas (HPA), the presence of reported epitopes extracted from literature and cancer-type recurrence in the dataset. Candidates were retained when their HPA tissue expression profile met at least one of the following criteria: no detected expression, expression restricted to placenta or testis, or expression below 10 nTPM across all tissues listed in the tissue-specific expression profile. Genes were further filtered based on the presence of at least one reported epitope.

### 2.13 Association between prioritized antigen gene expression and overall survival in bulk RNASeq data

Gene expression and overall survival analyses were performed using Bulk RNA-Seq data retrieved from the cBio-Portal for Cancer Genomics (Gao et al., 2013) through its REST API. For each cancer type, skin, liver, breast, pancreatic, colorectal, kidney, lung, ovarian, prostate, compatible TCGA studies were identified based on the availability of continuous mRNA expression profiles and clinical data. Gene expression values were used to stratify patients into high and low-expression groups based on the median expression among samples with values greater than 1, excluding zero-expression samples.

To avoid duplicate patient representation, one sample per patient was retained, and patients with conflicting expression classifications were excluded. Statistical significance was defined as p < 0.05, with survival associations classified according to the log-rank test.

### 2.14 Data Availability

All supplementary datasets generated during this study, including comprehensive gene lists, expression matrices, epitope validation results, and additional analytical outputs, are publicly available at Zenodo (https://zenodo.org/records/24431758)

## 3 Results

### 3.1 Dataset preprocessing and characterization

Originally, the dataset contained 355,941 cells, with the aim of improving quality control, cells annotated as Unknown were removed and T-cells were grouped. Unknown cells represented approximately 0.04%, 144 cells, leaving the dataset with 355,797 cells and group T-cell subtypes reduced the number of distinct cell types by nine, resulting in 21 cell types. Although the source dataset included cells from 10 tumor types, malignant cells were present in only nine: squamous cell carcinoma did not contain malignant cells, however, it was included in the analyses, except for those specifically focused on malignant cells (Figure 1A and Figure 2B).

**Figure 1:**
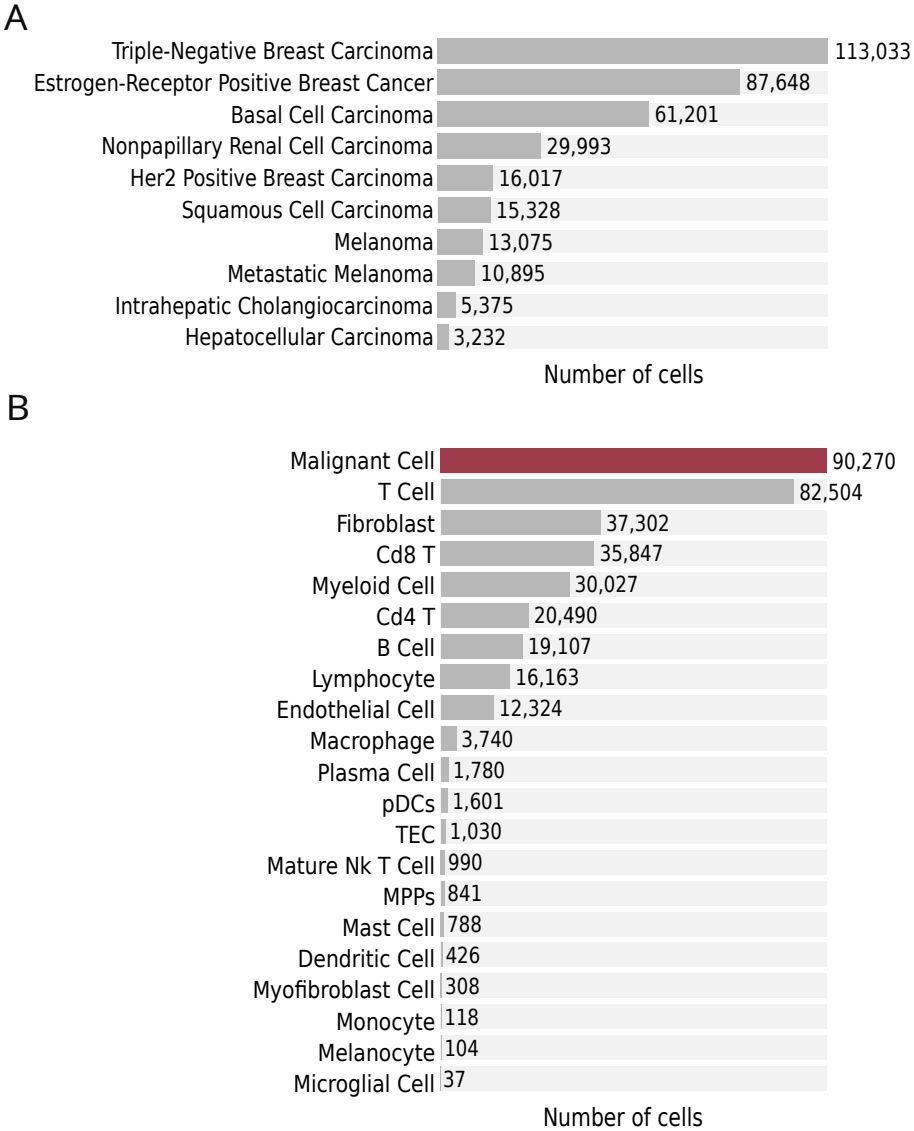
Composition of the integrated single-cell tumor dataset. (A) Number of cells across tumor types. (B) Number of cells across annotated cell types, including malignant cells (red) and non-malignant (gray) cells.

**Figure 2:**
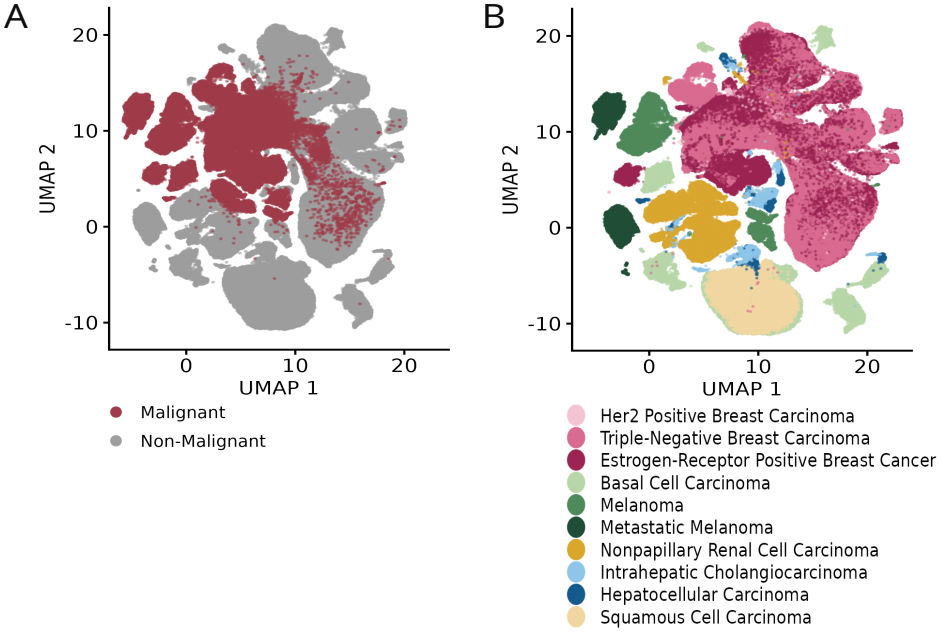
Composition of the integrated single-cell tumor dataset. (A) UMAP projection showing the distribution of malignant (red). (B) UMAP projection showing the distribution of cells according to tumor type, with each color representing a distinct tumor type. Each point represents a single cell. Abbreviations: UMAP, Uniform Manifold Approximation and Projection.

Malignant cells were the most abundant population, comprising 90,270 cells, approximately 25% of the dataset. T-cells are the second largest population, whereas microglial cells were the least represented, with 37 cells (Figure 1B and 2A). Triple-negative breast carcinoma contributed the largest number of cells, and hepatocellular carcinoma the smallest (Figure 1A).

### 3.2 A restricted set of genes is enriched in malignant cells across cancer types

All genes detected in the dataset were included in the cell-type specificity analysis, comprising 22,779 genes across 21 cell types. Most genes were classified as Low Specificity, while smaller proportions were assigned to the Group Enriched, Enhanced, or Enriched categories. In total, 226 genes were classified as Enriched in malignant cells and therefore considered candidates with malignant cell specificity within the scope of this study (Figure 3).

**Figure 3:**
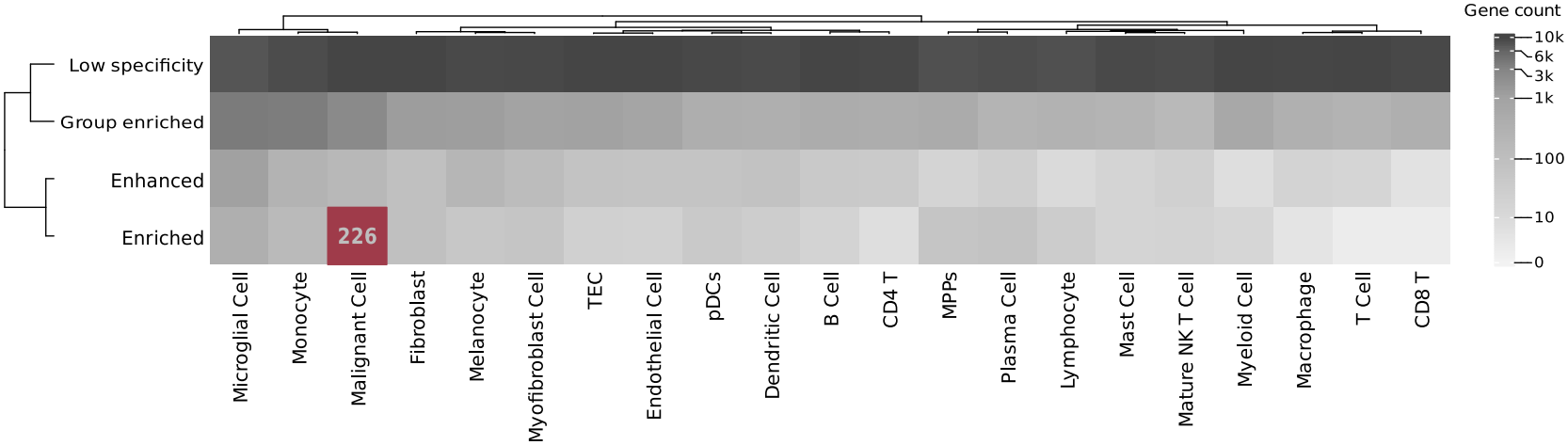
Distribution of cell-type specificity categories across annotated cell populations. Columns represent annotated cell populations, and rows represent specificity categories. Color intensity indicates the number of genes assigned to each category within each cell population, as shown by the gene count scale. Hierarchical clustering was applied to both dimensions, with dendrograms illustrating similarities in gene enrichment patterns across cell populations and specificity categories. The red annotation highlights the 226 genes classified as enriched in malignant cells. Abbreviations: TECs, thymic epithelial cells; pDCs, plasmacytoid dendritic cells; MPPs, multipotent progenitor cells.

### 3.3 Unexpected enrichment of cancer-testis antigens (CTAs) in non-malignant cells

Genes classified as Enriched, Enhanced, or Group Enriched were identified in non-malignant cell types, including CTAs reported as highly expressed in tumors (Naik et al., 2024). The highest CTA expression proportions and levels were observed in myeloid populations, with 25 CTAs associated with microglial cells and 26 with monocytes (Figure 4). Among the CTAs enriched in monocytes, those with the highest percentages of expressing cells were COX6B2 (43.22%), PIWIL2 (33.05%), and OTOA (27.96%). COX6B2 and PIWIL2 showed group enrichment in microglial cells and monocytes, whereas OTOA showed group enrichment in microglial cells, mono-cytes, and myeloid cells. ADAM29 (3.38%) and DMRT1 (3.38%) were classified as Enriched in monocytes.

**Figure 4:**
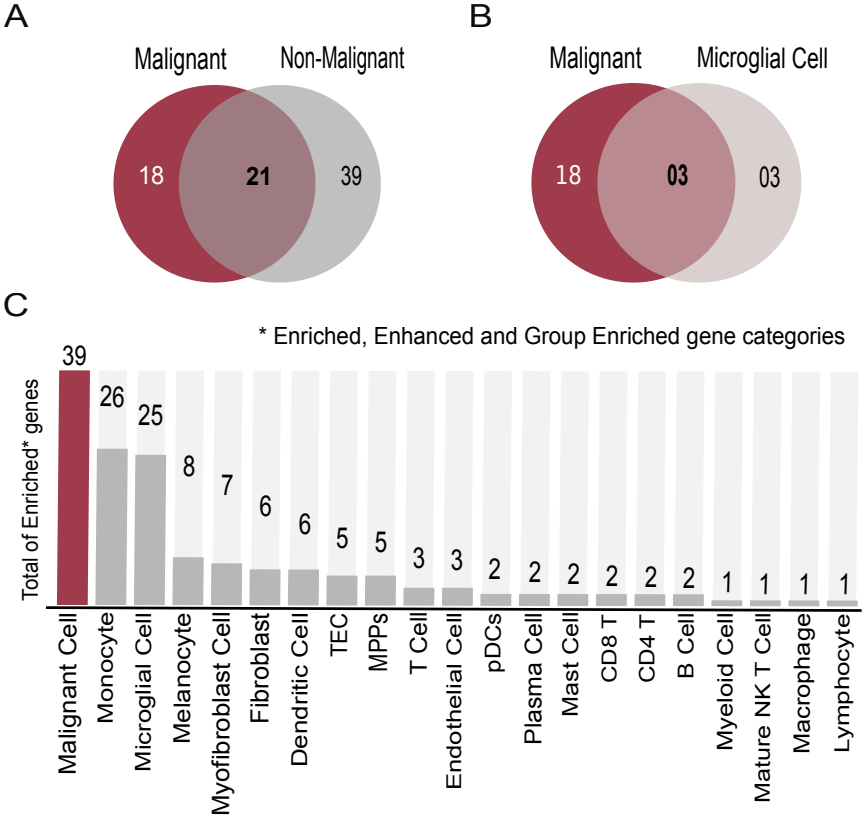
Distribution of enriched genes across malignant and non-malignant cell populations. (A) Venn diagram showing the overlap of genes classified as Enriched, Enhanced, or Group Enriched between malignant and non-malignant cell populations. Eighteen genes were exclusive to malignant cells, 39 were exclusive to non-malignant cells, and 21 were shared between both populations. (B) Venn diagram comparing malignant cells and microglial cells, showing 18 genes exclusive to malignant cells, three shared genes, and three exclusive to microglial cells. (C) Number of genes classified as Enriched, Enhanced, or Group Enriched across cell types. Malignant cells presented the highest number of genes (n = 39), followed by monocytes (n = 26) and microglial cells (n = 25).

In microglial cells, the most abundant CTAs were COX6B2 (48.65%), classified as group enriched in microglial cells and monocytes; ACRBP (43.24%), classified as enriched; PIWIL2 (37.83%), classified as group enriched in microglial cells and monocytes; and HSPB9 (16.21%), classified as enriched. In fibroblasts, the most abundant CTAs were SPEF2 (4.39%), classified as group enriched in dendritic cells, malignant cells, and fibroblasts; and ELOVL4 (4.14%), classified as group enriched in fibroblasts, myofibroblast cells and CD8 T. For the complete list of CTAs in this category, see the Supplementay data.

To further investigate the expression of selected CTAs in non-malignant tissues, single-cell RNA-sequencing data from normal tissues collected from HPA were analyzed. Among the selected genes, COX6B2 (CT59) showed expression in 0.1 nTPM of microglial cells and 0.2 nTPM of monocytes. PIWIL2 was detected in 5.5 nTPM of microglial cells and 0 nTPM of monocytes. OTOA was expressed in 7.6 nTPM of microglial cells and 0.1 nTPM of monocytes. ACRBP (CT13/OY-TES-1) was detected in 5.7 nTPM of microglial cells. Regarding fibroblasts, SPEF2 was expressed in 55.8 nTPM of these cells, with no additional information identified for other cell populations. Similarly, ELOVL4 was detected in 13.5 nTPM of fibroblasts and 3.9 nTPM of CD8 T cells. For the complete list of CTAs in this category, see the Supplementary data.

### 3.4 A subset of candidate antigens has epitopes reported in the literature

The 226 candidate antigen genes were assessed for previously reported epitopes in the databases searched. Of these, 113 had at least one reported epitope (Figure 5A), and some were associated with multiple epitopes. Of the 113 genes, 25 were classified as long intergenic non-coding RNAs (LINCs), while 18 corresponded to other non-coding RNA categories. Among the four broad cancer groups represented in the source dataset (breast, liver, kidney, and skin), epitopes were reported for breast and skin cancers. Epitope associations with lung, colorectal, ovarian, pancreatic, and prostate cancers were also identified (Figure 5B).

**Figure 5:**
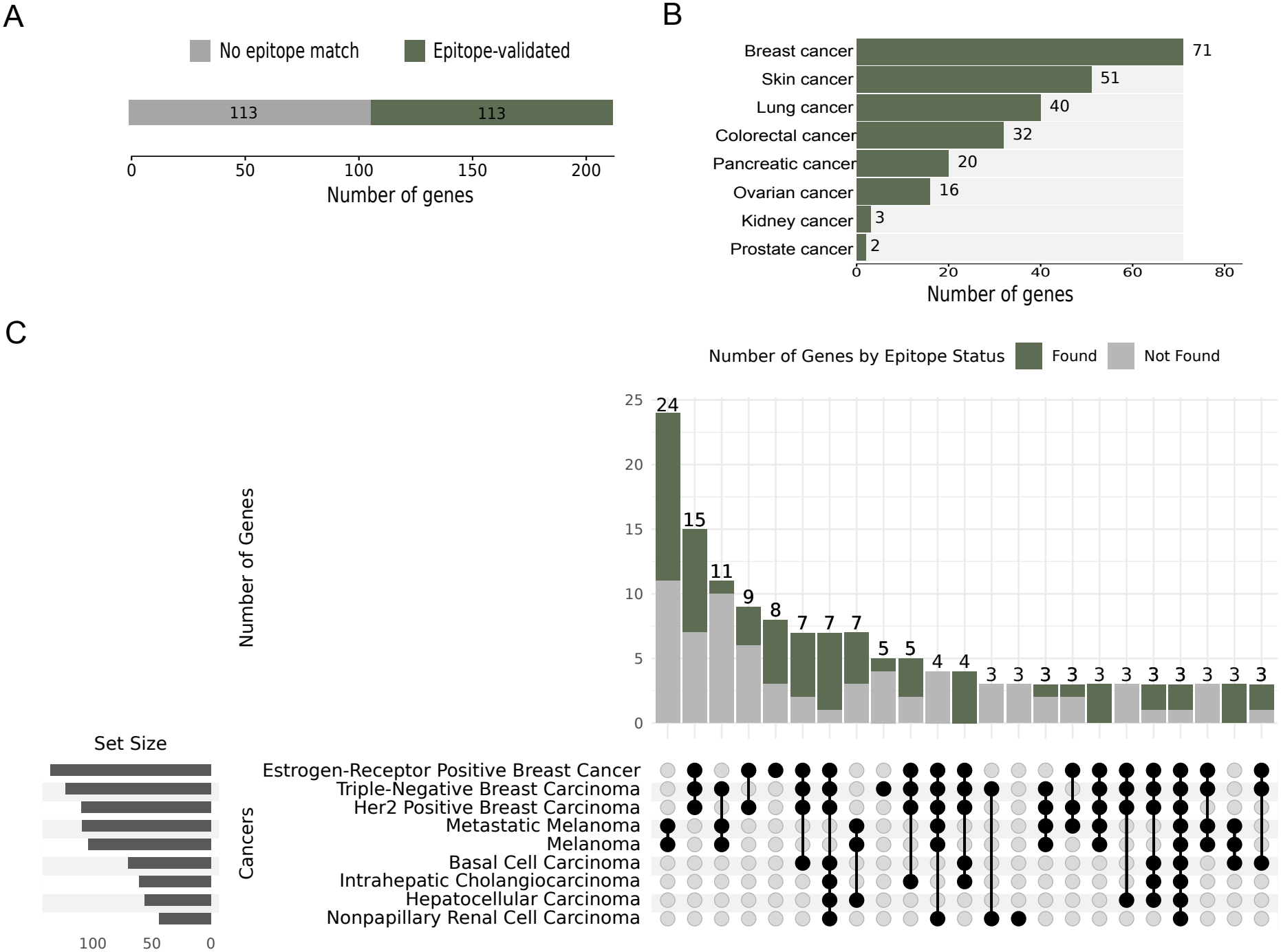
Overlap of epitope-associated genes across cancer types. (A) Number of candidate antigen genes with and without a reported epitope association based on epitope annotation databases. Genes classified as epitope-validated have at least one reported epitope association, whereas genes classified as no epitope match lack a corresponding epitope record. (B) Number of epitope-associated genes according to the cancer types for which epitopes have been reported in the public databases. Bar lengths and labels indicate the number of genes associated with each cancer type. (C) The UpSet plot summarizes the distribution and intersections of genes with reported epitopes among different cancer types. Horizontal bars on the left (Set Size) indicate the total number of epitope-associated genes reported for each cancer type. Vertical bars at the top represent the number of genes shared among specific combinations of cancer types. Connected black dots identify the cancer types included in each intersection, whereas unconnected dots indicate cancer types not included in that combination. Dark gray segments represent genes with reported epitopes (Found), and light gray segments represent genes without reported epitopes (Not Found).

### 3.5 Identification of candidate antigen genes shared across cancers

The largest pairwise overlap was between melanoma and metastatic melanoma, with 24 shared genes and the second one are the three breast cancer studies: estrogen-receptor positive breast cancer, triple-negative breast carcinoma and HER2 positive breast carcinoma (Figure 5C). CYP2J2, LINC00240, and MYO5B were detected in malignant cells from all nine tumor types. LINC00240 is a long non-coding RNA (lncRNA).

### 3.6 Immunotherapy-relevant candidate antigen genes retrieved from the literature

Three genes were selected for further discussion and highlighting because, in addition to being classified as enriched exclusively in malignant cells in the dataset and having documented epitopes in the literature, and are being investigated in the context of at least one immunotherapy modality.

#### High expression vs. Survival in bulk RNASeq data

As shown in (Figure 6), Kaplan-Meier survival analyses revealed significant associations between gene expression levels and overall survival across multiple cancer types. High expression of genes such as FLRT1 in breast cancer, CPA6 in colorectal cancer, and IL20 in breast cancer was associated with poorer survival outcomes, suggesting a potential role in tumor progression. Conversely, elevated expression of HOXC13 and ITIH6 in skin cancer was associated with improved survival, indicating possible protective or immune-related functions. These results highlight that the prognostic impact of individual genes is highly cancer-type specific and may provide insights into candidate for patient stratification and therapeutic targeting.

**Figure 6:**
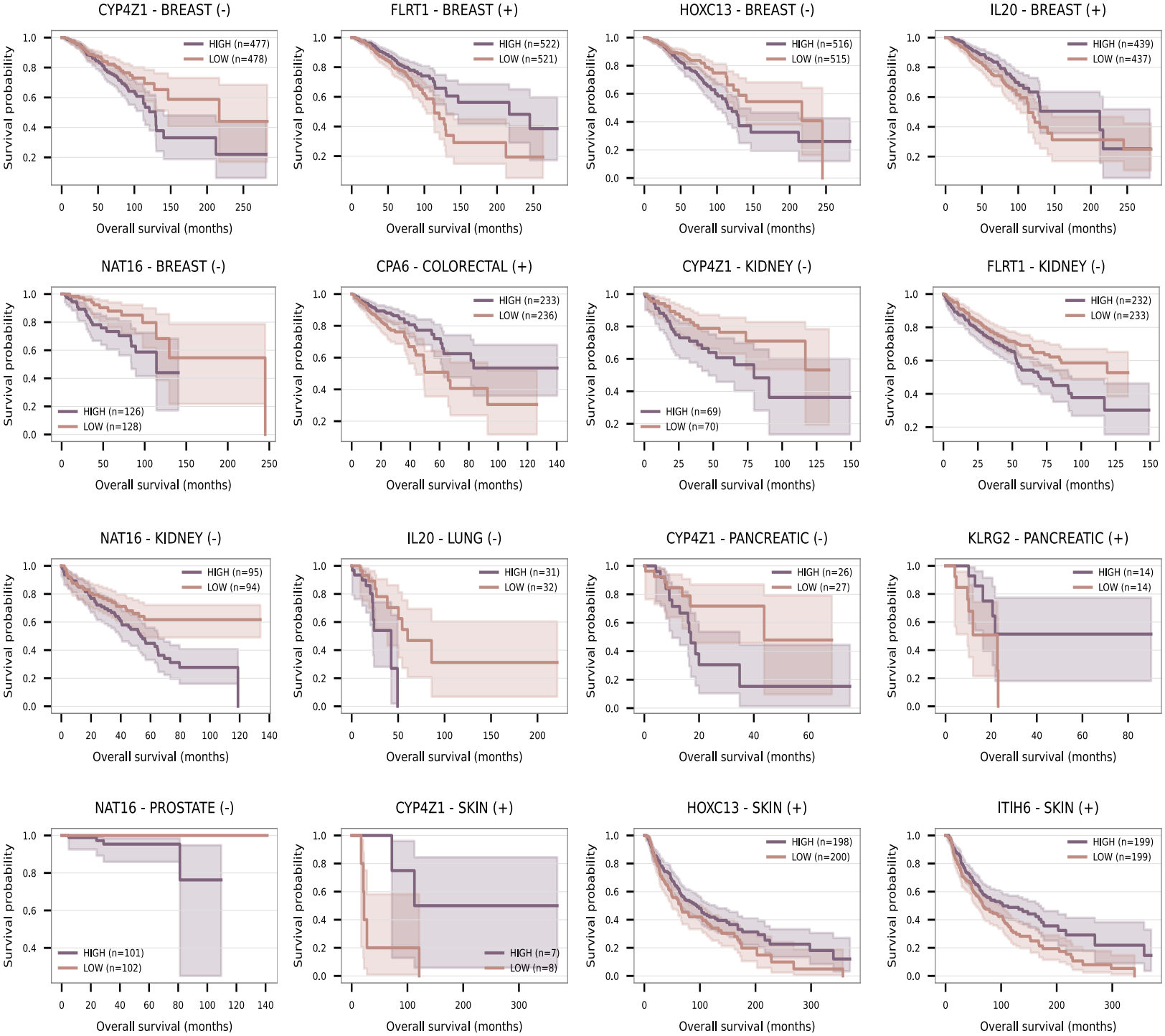
Overall survival analysis based on Kaplan-Meier curves for genes differentially associated with prognosis across multiple cancer types. Comparison between patients with high (HIGH) and low (LOW) gene expression levels allowed the identification of positive (+) or negative (-) associations with survival outcomes. Genes associated with improved prognosis may reflect biological mechanisms involved in antitumor immune responses, whereas genes associated with poorer prognosis represent potential therapeutic targets for molecular inhibition or targeted immunotherapy approaches. Shaded areas indicate the confidence intervals of the survival estimates.

## 4 Discussion

With the advancement of cancer immunotherapy strategies, the identification and characterization of potential tumor antigens have received increasing attention in scientific research. In this context, transcriptomic approaches have been widely employed to identify genes associated with malignancy and prioritize potential tumor antigens (Scurr et al., 2020; Yi et al., 2024; Schreiner et al., 2021), including in previous studies conducted by our research group (Fonseca et al., 2016). However, some transcriptomic approaches do not incorporate cellular heterogeneity into their analyses or consider non-malignant populations as a single reference group (Yang et al., 2023; Saul; Kosinsky, 2021). This approach may fail to capture the full extent of cellular heterogeneity, as gene expression in specific nonmalignant populations may be masked when these cells are analyzed collectively. To obtain a broader perspective on potential tumor antigen candidates derived from transcriptomic data, considering enrichment classifications across distinct cell populations within the tumor microenvironment provides additional information, representing an initial step toward prioritizing potential candidate antigen genes.

Beyond the identification of potential tumor antigen candidates, investigating the relative enrichment of genes and understanding the heterogeneity of malignant and non-malignant cells also allows the assessment of the expression patterns of classical antigens, such as CTAs. CTAs are a group of genes whose expression is predominantly observed in non-somatic tissues, particularly germline tissues, and aberrantly expressed in cancer. In this study, contrary to expectations, CTAs were not only detected in non-malignant populations but were also classified into enrichment categories in non-malignant populations. A total of 39 unique genes, suggesting that these genes are unsuitable targets for potential targeted immunotherapies, such as TCR-T cell therapy, TCR-based bispecific therapies, and cancer vaccines, as their targeting could result in unintended effects on non-malignant cells. Targeting a CTA enriched in non-malignant cells could affect normal cellular compartments and reduce therapeutic specificity.

Some of the CTAs classified as enriched in non-malignant cells also showed expression in normal tissues according to the datasets retrieved from the Human Protein Atlas (HPA), particularly SPEF2 and ELOVL4. Their detection in non-malignant cells in a tumor sample therefore does not necessarily indicate tumor-associated acquisition of expression. Conversely, candidates not detected in the evaluated healthy cell populations remain compatible with, but do not establish, a tumor-context association; further validation is needed.

Microglia and monocytes had the largest numbers of CTAs classified as enriched in non-malignant populations, despite being among the least represented cell types; microglia comprised only 37 cells. Because classification relied on the proportion of expressing cells within each cell type rather than absolute cell counts, differences in population size did not directly influence specificity assignments. Nevertheless, the small number of cells can reduce the precision of these estimates. Therefore, further studies involving larger numbers of cells and independent datasets are needed to investigate these specific CTAs and their association with expression and enrichment in non-malignant populations.

**Table 1:**
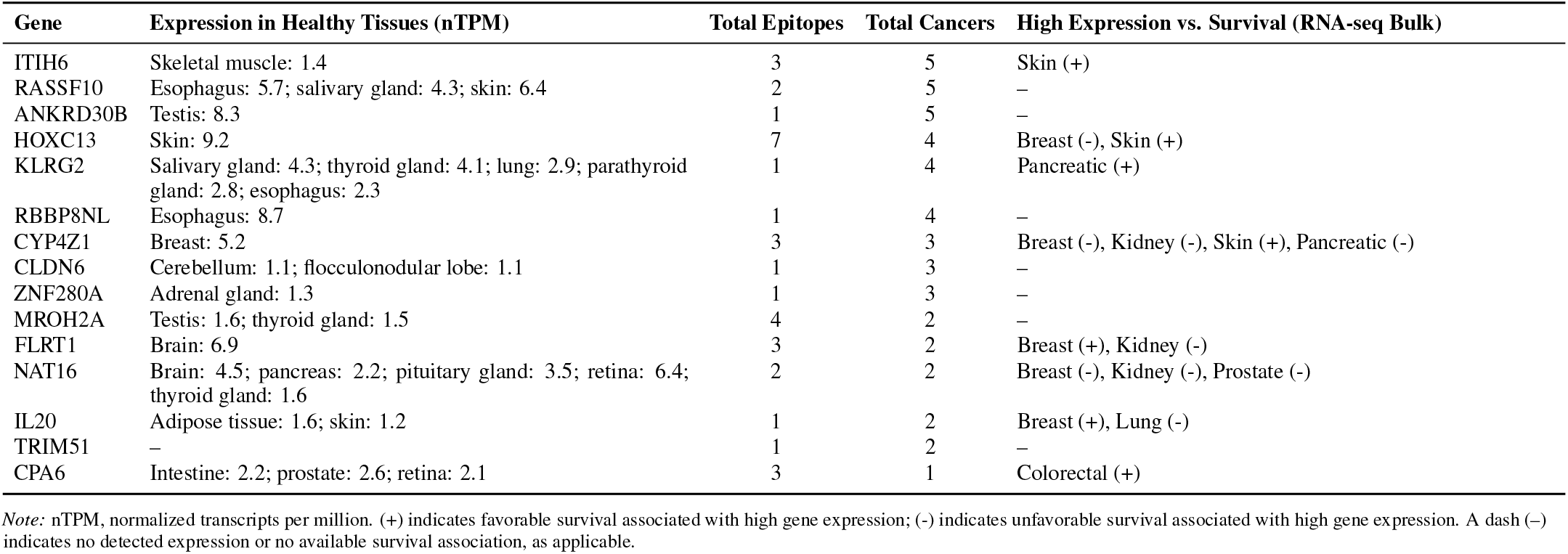
Characterization of prioritized candidate tumor antigens based on healthy tissue expression, epitope evidence, cancer recurrence, and survival analysis.

| Gene | Expression in Healthy Tissues (nTPM) | Total Epitopes | Total Cancers | High Expression vs. Survival (RNA-seq Bulk) |
|---|---|---|---|---|
| ITIH6 | Skeletal muscle: 1.4 | 3 | 5 | Skin (+) |
| RASSF10 | Esophagus: 5.7; salivary gland: 4.3; skin: 6.4 | 2 | 5 | – |
| ANKRD30B | Testis: 8.3 | 1 | 5 | – |
| HOXC13 | Skin: 9.2 | 7 | 4 | Breast (-), Skin (+) |
| KLRG2 | Salivary gland: 4.3; thyroid gland: 4.1; lung: 2.9; parathyroid gland: 2.8; esophagus: 2.3 | 1 | 4 | Pancreatic (+) |
| RBBP8NL | Esophagus: 8.7 | 1 | 4 | – |
| CYP4Z1 | Breast: 5.2 | 3 | 3 | Breast (-), Kidney (-), Skin (+), Pancreatic (-) |
| CLDN6 | Cerebellum: 1.1; flocculonodular lobe: 1.1 | 1 | 3 | – |
| ZNF280A | Adrenal gland: 1.3 | 1 | 3 | – |
| MROH2A | Testis: 1.6; thyroid gland: 1.5 | 4 | 2 | – |
| FLRT1 | Brain: 6.9 | 3 | 2 | Breast (+), Kidney (-) |
| NAT16 | Brain: 4.5; pancreas: 2.2; pituitary gland: 3.5; retina: 6.4; thyroid gland: 1.6 | 2 | 2 | Breast (-), Kidney (-), Prostate (-) |
| IL20 | Adipose tissue: 1.6; skin: 1.2 | 1 | 2 | Breast (+), Lung (-) |
| TRIM51 | – | 1 | 2 | – |
| CPA6 | Intestine: 2.2; prostate: 2.6; retina: 2.1 | 3 | 1 | Colorectal (+) |

*Note:* nTPM, normalized transcripts per million. (+) indicates favorable survival associated with high gene expression; (-) indicates unfavorable survival associated with high gene expression. A dash (–) indicates no detected expression or no available survival association, as applicable.

Mean expression across the dataset was highly skewed, with most genes showing low or undetectable expression and only a small fraction exhibiting high expression levels. Highly abundant transcripts included MALAT1, ACTB, EEF1A1, FTH1, FTL, mitochondrial genes, and CD74, a pattern commonly observed in single-cell RNA sequencing (scRNAseq) data. In contrast, the candidate tumor antigens identified in this study generally showed mean expression below 0.4. These findings indicate that absolute expression levels alone are insufficient for prioritizing therapeutic targets, supporting the use of enrichment-based classification and successive prioritization steps. The initial list comprised 226 genes enriched in malignant cells, which was reduced to 30 candidates after the prioritization process. These 30 genes, listed in the Supplementary data, were further investigated through a literature review to identify evidence of their use as targets in immunotherapies currently under investigation or in clinical practice. In addition, their expression was evaluated using previously published bulk RNAseq datasets to investigate potential associations between higher candidate gene expression and favorable or unfavorable prognosis across different cancer types, providing clinical context for the prioritization approach.

Three candidates were identified that are already being studied for potential targeted immunotherapies: CLDN6, MAGEC2 e PLAC1. Their supporting evidence was heterogeneous. MAGEC2, PLAC1, and CLDN6 combined evidence of immune recognition or therapeutic development with limited expression in the normal tissues evaluated. MAGEC2 recurred in four tumor types (hepatocellular carcinoma, intrahepatic cholangiocarcinoma, melanoma, and metastatic melanoma) and had the largest epitope count among the candidates, with 34 associations in breast, prostate, and skin cancers and MAGEC2 was tissue-enriched only in testis (18.4 nTPM) and had no equivalent expression in the other tissues evaluated, consistent with its CTA classification. MAGEC2 is being investigated in study NCT04729543 in genetically modified T cells expressing a MAGE-C2/HLA-A2-specific T-cell receptor (TCR) for patients with advanced melanoma (ClinicalTrials.gov, 2021).

CLDN6, which was identified in basal cell carcinoma, intrahepatic cholangiocarcinoma, and triple-negative breast carcinoma, has been evaluated using CAR-T cells alone or with an RNA vaccine in the BNT211-01 study (NCT04503278) (Mackensen et al., 2023). CLDN6 had very low expression in the evaluated normal tissues, with 1.1 nTPM in cerebellum and flocculonodular lobe. A response rate of 33% was reported among 21 evaluable patients. By contrast, the study of the bispecific antibody ASP1893 (NCT06681870) was withdrawn before initiation and provides no evidence of clinical activity (Clinical-Trials.gov, 2024). MAGEC2 and CLDN6 therefore have therapeutic strategies that have reached clinical evaluation, although the evidence does not by itself establish efficacy for the tumor contexts in which they were identified here. PLAC1 was enriched only in estrogen receptor-positive breast cancer and had three epitope associations with breast and colorectal cancers. In normal tissues, its expression was primarily restricted to the placenta (26.2 nTPM). A 2025 study evaluated a PLAC1-targeted antibody-drug conjugate in a murine breast cancer model, providing preclinical evidence of therapeutic potential (Jafar Mahmoudian et al., 2025).

In addition to these three genes, 27 other genes met the prioritization criteria. Among them, 15 do not fall into the classical categories of tumor antigens, such as cancertestis antigens (CTAs), overexpressed antigens, or differentiation antigens. Nevertheless, some have already been identified by the Human Protein Atlas (HPA) as potential prognostic markers, associated with either favorable or unfavorable outcomes. These include RASSF10 and KLRG2, which are associated with favorable prognosis in lung adenocarcinoma, and RBBP8NL, which is associated with favorable prognosis in bladder urothelial carcinoma. HOXC13 was associated with unfavorable prognosis in glioblastoma, whereas FLRT1 showed a favorable association. NAT16 was associated with favorable prognosis in cutaneous melanoma but unfavorable prognosis in thyroid carcinoma.

Survival analyses revealed distinct prognostic associations among the candidate genes, suggesting potential differences in their biological roles across cancer types. Genes whose higher expression was associated with better survival, including ITIH6 (skin), KLRG2 (pancreatic), CPA6 (colorectal), FLT1 (breast), HOXC13 (skin), and CYP4Z1 (skin), may reflect biological processes associated with less aggressive tumor phenotypes or a more favorable tumor microenvironment. Conversely, genes whose higher expression was associated with poorer survival, including HOXC13 (breast), CYP4Z1 (breast, kidney, and pancreatic), NAT16 (breast, kidney, and prostate), IL20 (breast and lung), and FLT1 (kidney), may represent candidates for further investigation as therapeutic targets. Depending on their protein localization and biological relevance, potential approaches could include monoclonal antibodies, small-molecule inhibitors, or RNA-based strategies. Furthermore, regardless of their prognostic associations, genes encoding intracellular proteins may also represent potential immunotherapeutic targets if their derived peptides are presented by HLA molecules and recognized by tumor-specific T cells. These targets could be explored through therapeutic vaccines or T-cell therapies involving lymphocytes engineered to express specific T-cell receptors (TCRs), whereas conventional CAR-T approaches would require the presence of antigens accessible on the cell surface.

As future research directions, cell-cell communication analyses could investigate potential interactions between malignant cells and immune populations within the tumor microenvironment, considering the expression of candidate genes and their respective ligands and receptors. This approach could help identify potential mechanisms of immune response modulation and clarify whether certain candidates are associated with interactions that promote or impair tumor cell recognition. Additionally, investigating the subcellular localization and protein expression of the candidates using proteomic data would be important to assess their accessibility as therapeutic targets and identify those with potential for CAR-T therapy development. Finally, spatial transcriptomics analyses could help characterize the distribution of candidate genes within tumor tissues and their proximity to different immune cell populations, while studies of intratumoral heterogeneity could determine whether their expression is distributed across most malignant cells or restricted to specific subpopulations.

## 5 Conclusion

Overall, this study provides an integrative framework for prioritizing candidate tumor antigens across multiple cancer types by combining single-cell transcriptomic profiles, healthy tissue expression, immunological evidence, and clinical associations. The identification of 30 prioritized candidates, alongside 39 cancer-testis antigens enriched in non-malignant populations, highlights the importance of considering cellular heterogeneity and potential off-tumor expression when assessing therapeutic suitability. While some candidates, including CLDN6, MAGEC2, and PLAC1, have already attracted therapeutic investigation, the findings reinforce that malignant-cell enrichment alone is insufficient to establish target safety or clinical relevance. Further characterization, including protein-level validation, HLA-restricted antigen presentation, subcellular localization and spatial expression, is necessary to establish their therapeutic potential. Ultimately, this integrative approach provides a foundation for the further investigation of candidate tumor antigens and can support the development of more selective cancer immunotherapies.

## Acknowledgments

The authors would like to thank the Graduate Program in Bioinformatics at the Federal University of Rio Grande do Norte (UFRN) for the academic support and infrastructure provided during the development of this study. We also acknowledge the contributions of colleagues and collaborators whose discussions and support helped improve this work.

